# Inflamed Natural Killer cells with adhesion defects are associated with a poor prognosis in Multiple Myeloma

**DOI:** 10.1101/2024.01.15.575654

**Authors:** Eve Blanquart, Rüçhan Ekren, Bineta Rigaud, Marie-Véronique Joubert, Virginie Baylot, Hélène Daunes, Marine Cuisinier, Marine Villard, Nadège Carrié, Céline Mazzotti, Virginie Baylot, Liliana E. Lucca, Aurore Perrot, Jill Corre, Thierry Walzer, Hervé Avet-Loiseau, Pierre-Paul Axisa, Ludovic Martinet

**Author notes:** Equal first authors (Eve Blanquart, Rüçhan Ekren, Bineta Rigaud). Equal senior authors (Pierre-Paul Axisa, Ludovic Martinet). **Lead contact**: Dr Ludovic Martinet, INSERM UMR 1037, Cancer Research Center of Toulouse, 2 av Hubert Curien, 31037 Toulouse, France. E-mail address; For original data, please contact.

## Abstract

The promising results obtained with immunotherapeutic approaches for multiple myeloma (MM) call for a better stratification of patients based on immune components. The most pressing being cytotoxic lymphocytes such as Natural Killer (NK) cells that are mandatory for MM surveillance and therapy. In this study, we performed a single cell RNA sequencing analysis of NK cells from 10 MM patients and 10 age/sex matched healthy donors (HD) that revealed important transcriptomic changes in NK cell landscape affecting both the bone marrow and peripheral blood compartment. The frequency of mature cytotoxic “CD56^dim^” NK cell subsets was reduced in MM patients at the advantage of late-stage NK cell subsets expressing NFκB and IFN-I inflammatory signatures. These NK cell subsets accumulating in MM patients were characterized by a low CD16 and CD226 expression and poor cytotoxic functions. MM CD16/CD226^lo^ NK cells also had adhesion defects with reduced LFA-1 integrin activation and actin polymerization that may account for their limited effector functions *in vitro*. Finally, analysis of BM infiltrating NK cells in a retrospective cohort of 177 MM patients from the IFM 2009 trial demonstrated that a high frequency of NK cells and their low CD16 and CD226 expression were associated with a shorter overall survival. Thus, CD16/CD226^lo^ NK cells with reduced effector functions accumulate along MM development and negatively impact patients’ clinical outcome. Given the growing interest in harnessing NK cells to treat myeloma, this improved knowledge around MM-associated NK cell dysfunction will stimulate the development of more efficient immunotherapeutic drugs against MM.

**Scientific category:** Lymphoid Neoplasia; Immunobiology and Immunotherapy.

**KEY POINTS:**

- MM patients have increased CD16/CD226^low^ NK cell subsets characterized by “inflammatory” signatures and reduced effector functions.
- The frequency of CD16/CD226^low^ NK cells correlate with MM patient clinical outcome

## Introduction

Multiple myeloma (MM) is an incurable hematological malignancy characterized by the proliferation of clonal, long-lived plasma cells (PCs) within the bone marrow (BM) associated with bone destruction, serum monoclonal gammopathy and organ dysfunction. The introduction of proteasome inhibitors (PI), immunomodulatory drug (IMIDs) and more recently monoclonal antibodies (mAbs) targeting CD38 has represented a major breakthrough in the treatment of newly diagnosed MM (NDMM) ^1,2^. In addition, Bispecific antibodies and chimeric antigen receptor (CAR) T cells targeting BCMA or GPRC5D on malignant PC have shown important therapeutic promise against relapsed/refractory MM (RRMM) suggesting that immunotherapy represents the future of MM treatment ^3,4^. Yet, most patients experience recurrent relapses with reduced therapeutic options which call for a better understanding of the resistance mechanisms to immunotherapy. While genetic alterations within malignant PC and the presence of aggressive sub-clones play a critical role in MM clinical outcome ^5,6^, recent studies also points toward the importance of the immune microenvironment in the pathogenesis of this disease ^7–9^. Indeed, myeloma development is associated with progressive immune dysfunctions that contribute to myeloma growth and drug-resistance. These include notably, dysregulated production of IL-18 that drives MDSC immunosuppression in the MM microenvironment ^10^, and CD8^+^ T dysfunction due to an imbalance between TIGIT inhibitory and CD226 (DNAM-1) activating receptors ^11,12^.

Natural Killer (NK) cells are innate cytotoxic lymphocytes that efficiently kill autologous MM cells *in vitro* ^13,14^ and are required for myeloma immunosurveillance *in vivo* ^7^. NK cells also kill antibody-coated target cells through CD16 (FcgRIII), a process called antibody-dependent cellular cytotoxicity (ADCC), and these cells are critically involved in the anti-myeloma efficacy of anti-CD38 mAbs ^15–17^. Finally, NK cells can acquire memory-like properties, such as long-term persistence and enhanced functions making these cells attractive therapeutic targets against myeloma ^18^.Thus, accumulating evidence suggest the existence of an NK cell-mediated pressure in MM and there is a growing interest around their therapeutic manipulation ^19–21^. Yet, key phenotypic and functional assessments of NK cells in MM patients are missing, with several articles reporting conflictive results.

In this study, through single cell RNAseq, spectral flow cytometry and functional analysis on MM patients’ samples, we found that NK cells characterized by a low CD16 and CD226 expression, inflammatory signatures, adhesion defects and reduced effector functions negatively impact MM patients clinical outcome.

## METHODS

### Human samples

Healthy donor PBMC were obtained from the Etablissement Français du Sang (EFS). BM aspirates from healthy donors were purchased from cliniscience and Lonza. Fresh BM aspirates and PB from patients with MM were collected at the time of diagnosis or relapse in the Institut Universitaire du Cancer de Toulouse-Oncopole (IUCT-O). All donors and patients gave their written informed consent and collection was approved by French Committee for the Protection of Persons (CPP; DC-2012-1654) and by local IUCT-Oncopole review boards.

### Cell lines and cell culture

P815 Mastocytoma (ATCC TIB-64) and K562 (ATCC CCL-243) cell lines were grown in complete DMEM medium containing 10% heat-inactivated fetal calf serum (FCS) and were tested negatively for mycoplasma contamination.

### Human NK cell functional assays

NK cells were purified using NK Cell Isolation Kit (Miltenyi Biotech) or FACS melody™ (BD Biosciences, US) and were kept overnight in complete RPMI media with 50 ng/ml of recombinant human IL-2 (R&D Systems). NK cells from HD or MM patients were stimulated in 96 wells plates with plate bound anti-CD16 (clone 3G8 Biolegend, 10ug/ml) or with IL-12 (Thermo Fisher Scientific, 10ng/ml), IL-15 (R&D Systems, 10ng/ml), IL-18 (Thermo Fisher Scientific, 20ng/ml), Phorboll 12-myristate 13-acetate (50ng/ml, Sigma-Aldrich) and ionomycine (1ug/ml, Sigma-Aldrich), alone or in combination. NK cells were also co-cultured with K562 or P815 coated with anti-NCR (NKp30, NKp44, NKp46, R&D Systems, 5ug/ml) or anti-CD16 (clone 3G8, 5ug/ml) or irrelevant IgG1 (clone MG1-45, Biolegend, 5ug/ml) at a 5:1 effector: target ratio.

### NK cell conjugates

PB NK cells from HD or MM, stained with cell trace violet (Thermofisher scientific, 5µM), were incubated with cell trace far red (Thermofisher scientific, 5µM) labelled K562 at a ratio of 1:1 for 1 hour and conjugate formation was evaluated by flow cytometry (Fortessa X20™, BD Biosciences).

### NK cell imaging

NK cells were incubated on recombinant Human ICAM-1-Fc (R&Dsystems) coated plates at 37°C for 20 minutes. Cells where fixed and permeabilized using Cytofix/Cytoperm™ Fixation/Permeabilization Kit (BD biosciences), the polymerized actine was stained using Phalloidin (Abcam) and LFA-1 activation was evaluated using m24 antibody (BioLegend). Alternatively, NK cells were incubated on poly-Lysine coated plates in the presence of Acridine Orange/Propidium Iodide Stain (Logos Biosystems, 20µg/ml) and imaged for 30 minutes with one picture taken every minute. Images were taken on an Operetta CLS analysis system (PerkinElmer) and analysis was done using the Harmony software.

### Flow cytometry

Cells were stained with fixable viability dye for 10 minutes in the dark. Extracellular staining was done in FACS Buffer (2% SVF, 5mM EDTA), with True stain Monocyte blocker™ (Biolegend) and BD Horizon™ Brilliant Stain Buffer Plus (BD Biosciences) following manufacturer instructions. For intracellular staining the Cytofix/Cytoperm™ Fixation/Permeabilization Kit (BD biosciences) was used following manufacture instructions. Acquisition was performed on a Fortessa™ X20 (BD Biosciences) or an Aurora (Cytek®). Data Analysis was performed using FlowJo or OMIQ softwares.

### Single cell RNA sequencing

BM and PB Cell suspensions were stained with Fixable Viability Dye eFluor 780 (Invitrogen), anti-CD3 (UCHT1), anti-CD56 (NCAM16.2), anti-CD138 (MI15) (all from BD Biosciences), and oligonucleotides-barcoded antibodies (TotalSeq™-B, Biolegend). Live NK (CD3-CD56+CD138-) and T cells (CD3+CD138-) from each donor were FACS sorted (ARIA II Fusion, BD Biosciences) and mixed at a 1:2 ratio of NK:T cells. Beads-cells emulsions were prepared using the Chromium Single Cell 3’ kit (10x genomics) per manufacturer’s instructions. Single-cell library size and quality were confirmed on the Fragment Analyzer (Agilent). KAPA Quantification Kit for Illumina platforms (KAPA Biosystems, Roche) was used to quantify libraries. The libraries were sequenced on a NextSeq 550 (Illumina) in pair-end sequencing 28 bp (read1) × 91 bp (read2) and a single index 8 bp in length.

### Survival analysis

Overall survival was calculated from MM diagnosis to death from any cause. Kaplan-Meier curves and p-values from log-rank were computed using the survival package in R (version 2.38). Cox proportional hazard models were adjusted using the survival package in R (Version 3.5-7; https://github.com/therneau/survival) and effects tested using ANOVA from the limma package.

### Statistics

Statistical analyses were performed using GraphPad Prism 9 Software. Mann-Whitney *U* test or unpaired student *t* test was used for single comparisons between two groups. For comparison of three or more groups, one-way ANOVA with Tukey’s multiple comparison test, Holm-Sidak multiple test correction, or non-parametric Kruskal-Wallis test with Dunn’s multiple comparison post-test were used.

## RESULTS

### Alteration of NK cell transcriptomic landscape in MM patients

To better understand the NK cell landscape in MM patients, we performed a CITE-Seq (Cellular Indexing of Transcriptomes and Epitopes by Sequencing) analysis of NK cells with the 10x Genomics high-throughput droplet-based scRNAseq pipeline. Paired blood and BM NK and T cells, defined respectively as CD3^−^CD56^+^ and CD3^+^ T cells were sorted by flow cytometry from 10 MM patients and 10 age/sex matched healthy donors (HD). After filtering cells using standard quality controls, an *in silico* extraction of cells expressing NK public cell signatures was performed (**Figure S1A-B and supplemental method section**). This automated approach was manually validated by gene and antibody derived tag (ADT) expression analysis (**Figure S1C-D**). A total of 22,749 BM and 14,182 blood NK cells were clustered into eight distinct clusters (C1-C8) without detectable contaminants with other cell lineages (**Figure 1A)**. Following integration (**supplemental method section**), the different HD and MM NK cells from BM and PB were evenly distributed in the different clusters (**Figure S1E-G)**. Analysis of PB and BM NK cell landscape, through cluster frequency analysis or MELD likelihood algorithm (**supplemental method section**), revealed important differences between MM patients and HD (**Figure 1B-C**). The frequencies of cluster 1 and 2, the most abundant NK clusters in PB and BM from HD samples, were reduced in MM patients (**Figure 1B-C**). These clusters had signatures (*Crinier et al. 2018* ^22^*, Yang et al. 2019* ^23^) of CD56^dim^ mature NK cells and were characterized by high expression of genes related to cytotoxicity (PRF1, GZMA, GZMB, CD226 and FCGR3A) (**Figure 1D-E and Figure S1H**). Conversely, MM patients had a significant increase in cluster 3 frequency that represented the majority of BM and PB MM NK cells while they only accounted for a small fraction of HD NK cells (**Figure 1B-C**). This cluster was characterized by “activated/inflammatory” CD56^dim^ NK cell signatures and had high expression of NFκB related genes (NFKB1, REL, RELB) (**Figure 1D-E**). A population of NK cells (cluster 4) characterized by adaptive-like NK cell signature genes (KLRC2, KLRC3, CD3E and IL32) was found in the BM and PB of HD (**Figure 1D-E**). Interestingly, this cluster of “adaptive-like” NK cells was reduced in MM patients at the expense of a second cluster of NK cells with adaptive-like features (**cluster 7**) that expressed type 1 interferon (IFN-I) related genes (IFIT1, IFIT2, IFIT3, IFIH1) (**Figure 1B-E**). Both MM associated clusters C3 and 7 were characterized by a lower expression of FCGR3A (encoding CD16) (**Figure 1E and S1H**). Of note, we did not identify important differences between BM and PB, except cluster 8 that was mainly restricted to BM samples and characterized by the expression of BM resident NK cells (BMrNK) signatures ^22–24^ (**Figure 1D-E and Figure S1H-I).** The frequencies of this cluster as well as cluster 5 and Cluster 6 characterized by CD56^bright^ and terminal NK cell signatures respectively (**Figure 1D-E**), were also similar between MM and HD, PB and BM NK samples (**Figure 1B**). Altogether these results show that MM is accompanied by important transcriptomic alterations of the BM and PB NK cell compartment with decreased mature cytotoxic CD56^dim^ NK cells and increased CD16^lo^ inflamed NK cell subsets.

**Figure 1:**
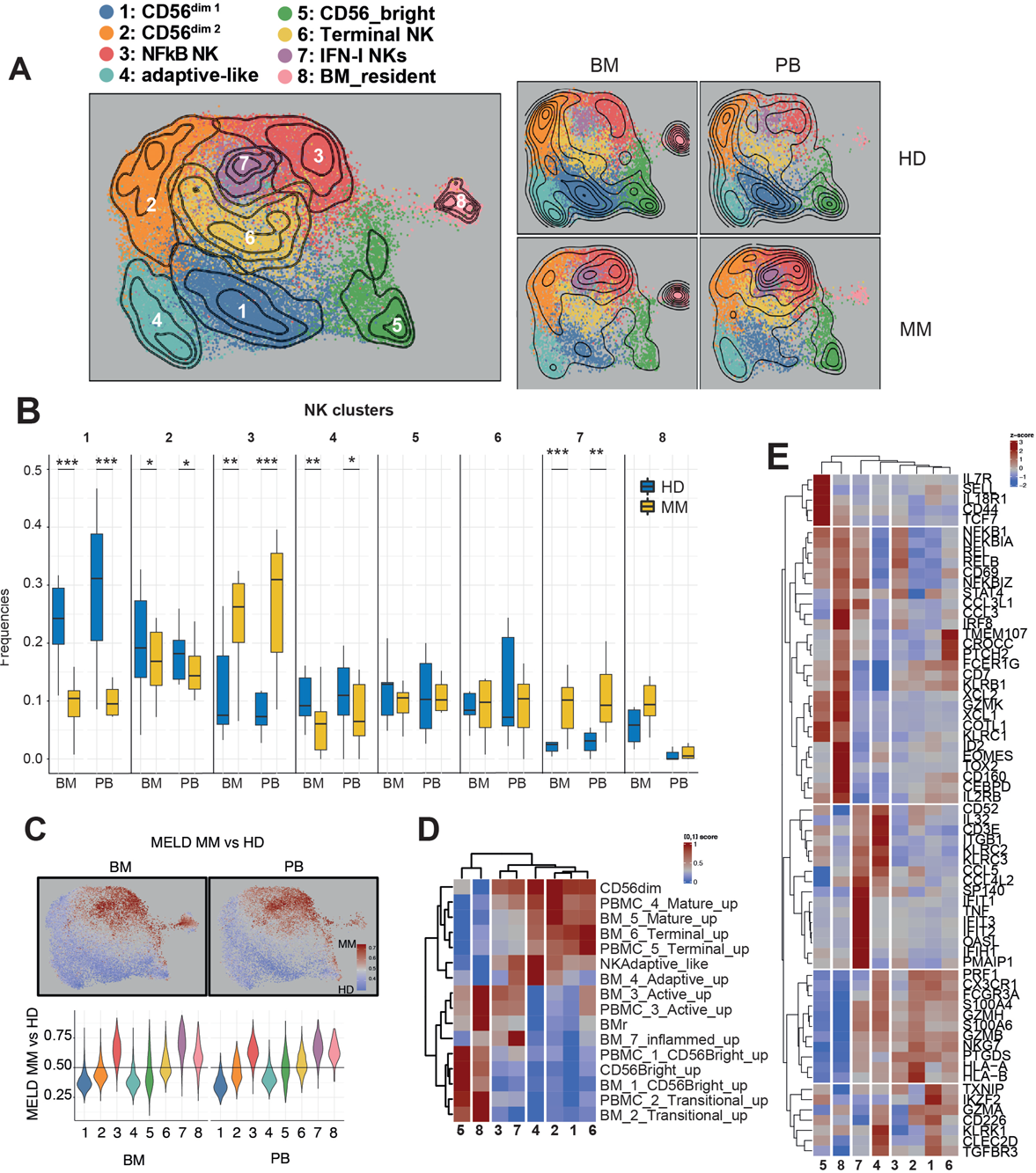
Alteration of NK cell transcriptomic landscape in MM patients. A scRNAseq analysis of Paired blood and BM CD3^−^CD56^+^ NK cells from 10 MM and 10 age/sex was performed. **A.** UMAP embeddings of NK cells colored by clusters. **B.** Variations of NK cluster frequency across disease and tissue status (box-and-whisker plots, where whiskers represent 1.5 * interquartile range and box represents the 25^th^, 50^th^, and 75^th^ percentiles; false discovery rate (FDR) derived from generalized linear model, see supplementary Methods; FDR < 0.05*, FDR < 0.01**, FDR < 0.001***). **C.** MELD likelihood (see supplementary Methods) highlighting MM vs HD enrichment computed separately on BM or PB. Likelihood values are overlayed on UMAP embeddings and per-cluster likelihood distributions displayed as violin plots. **D.** Heatmap of marker genes standardized expression across clusters. **E.** Heatmap of standardized gene module scores from published datasets (crinier et al. 2020 and Yang et al. 2019).

### Accumulation of late stage inflamed NK cells in Multiple myeloma

To better understand the developmental link between NK subsets differentially present between MM patients and HD, we performed a slingshot pseudotime trajectory analysis (**Figure 2A**). Based on the current human NK cell development model ^25^, we assigned the “CD56^bright^ NK” cluster 5 as the least mature branch in the pseudotime. BM and PB NK cells demonstrated a relatively simple developmental progression with few branches (**Figure 2B**). The “mature CD56^dim^” NK C1 and the “BMrNK” NK C8 followed two different developmental paths from CD56^bright^ NK cells (**Figure 2B-C**). “BMrNK” formed a unique separate branch (Lineage 3) while all the other clusters derived from the “mature CD56^dim1^” C1 cluster branch that gave rise to the “terminal NK C6” (Lineage 2) and the “mature CD56^dim2^” NK C2 (Lineage 1). The “adaptive-like” NK C4 emerged between “mature CD56^dim1^” and “mature CD56^dim2^” stages. Both “NFkB” C3 and “IFN-I NK” C7 derived from “mature CD56^dim2^” NK cluster C2 and dominated the end of the trajectory (**Figure 2B-C**). Analysis of MM vs HD NK cell distribution along pseudotime for the lineage 3 branch confirmed that MM patients’ NK cells were more prevalent than HD at latest stages of NK cell pseudotime in both PB and BM compartment (**Figure 2D**). Trajectory pseudotime correlated with MELD MM vs HD likelihood highlighting enrichment of MM cells at late pseudotime (**Figure 2E**). Of note, similar results were obtained when analysis cells from the bone marrow vs blood separately, and when using “BMr” NK C8 as a starting root (**Figure S2A-C**). Altogether, these results suggest that MM alters normal NK cell homeostasis pushing the cytotoxic CD56^dim^ NK cells C1 and adaptive NK subsets C4 to differentiate into “NFkB NK” C3 and “IFN-I NK” C7 (**Figure 2F**). To better appreciate the signal that contribute to this shift in NK cell development, we performed Gene set enrichment analysis (GSEA) between the NK cells cluster C1 vs C3 using the KEGG data base. We found that C3 NK cells had enriched expression of inflammation related pathways including “NOD like receptor”, “RIG1 like receptor” and “Toll like receptor” signaling pathways as compared to “CD56^dim”^ NK cells (**Figure 2G**). Similar enriched signatures of inflammation were found comparing adaptive-like NK clusters C7 vs C4 (**Figure 2H**) or comparing MM patients to HD NK cells (**Figure S2D**). Similar results were obtained using Reactome data base (**Figure S2E**). Leading edge genes for these pathways included several chemokines (CCL3, CCL5), TNF (TNFRI, TNF, TRAF3, RELA, MAP2K3, MAP2K7, MAP3K8, NFKB1, NFKBIA) and IL-1 family (TICAM1 TAB2 TBK1) related genes (**Figure 2I**). These mediators are often associated with MM development ^26,27^ and could be responsible for the late stage “inflamed NK cells” that accumulate in MM.

**Figure 2:**
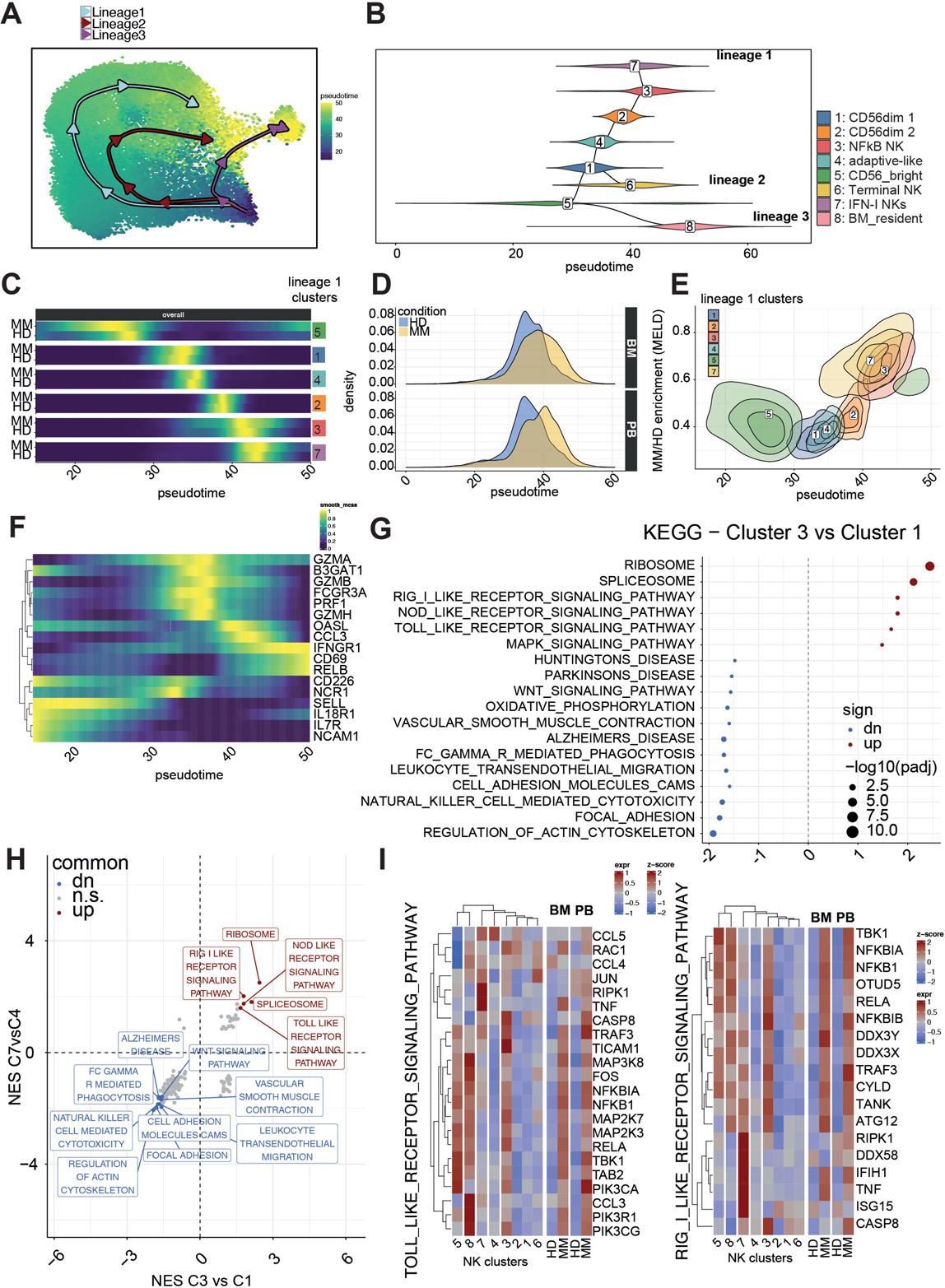
Accumulation of late stage inflamed NK cells in Multiple myeloma. **A.** Trajectory analysis results overlayed on UMAP embeddings identified 3 lineages seeding from CD56^bright^ NK cluster 5. **B.** Branching of the minimum spanning tree built at the cluster level differentiates bone marrow resident NKs, terminal NKs and inflammatory MM-enriched NK cells. **C.** Cell density distributions along trajectory pseudotime describes NK cells differentiation. **D.** Pseudotime distributions of PB and BM NK cells from MM patients and HD. **E.** Trajectory pseudotime correlation with MELD MM vs HD likelihood. **F.** Gene expression along pseudotime trajectory. **G.** Comparison of gene expression programs between cluster 3 and 1 using GSEA analysis of KEGG genesets. **H.** Comparisons between GSEA normalized enrichment scores (NES) of C7vsC4 and C3vsC1 NK clusters. **I**. Heatmaps of leading-edge genes standardized expression from the indicated inflammatory pathways across clusters and disease.

### Decreased cytotoxic functions in NK cell from MM patients

GSEA revealed that MM NK clusters C3 and C7 have decreased expression of cytotoxicity related signatures such as “Natural killer cell mediated cytotoxicity” and “FC gamma R mediated phagocytosis” (**Figure 2H**). Leading edge genes for these pathways included several activating receptors (FCGR3A, NCR1, CD244), key signaling molecules (SYK, LAT, LCK, VAV1, VAV3, ZAP70) and cytotoxic mediators (GZMB, PRF1) that were overall less expressed in MM than in HD NK cells (**Figure 3A)**. To validate these findings at the protein level, we next performed an unsupervised spectral flow-cytometry analysis of frozen BM samples from 21 HD and 49 NDMM (**Figure 3B and Figure S3A**). Despite important inter-individual heterogeneity, we confirmed that MM patients have increased frequency of NK cell subset characterized by low levels of cytotoxic mediators Perforin and granzyme B (**Figure 3C-F and Figure S3B**). Consistent with the phenotype observed by scRNAseq, these NK cells expressed low levels of CD226 and CD16 and had higher levels of CD69 (**Figure 3C-F**). Altogether, these results suggest limited effector functions in MM patients’ NK cells. We further explored this finding at the functional level by comparing PB NDMM and HD NK cell degranulation through CD107a and their IFNγ intracellular production in redirected lysis assay using mouse FcγR^+^ P815 cell line coated with agonist mAbs against CD16 or Natural cytotoxicity receptors (NCRs, NKp30, NKp44 and NKp46) (**Figure 3G-H**). We observed a significant increase in the expression of IFNγ and CD107a by HD NK cells in the presence of P815 coated with NCRs or CD16 mAbs as compared to Ig control. By contrast, the expression of IFNγ and CD107a was very low regardless of the condition tested for MM NK cells. Low MM NK cell reactivity was found upon stimulation with anti-CD16 coated wells or an IL-12/IL-15/IL-18 cytokine cocktail (**Figure S3C-D)**. Even PMA/Ionomycin only induced a low increase in IFNγ production by MM NK cells while it strongly stimulated HD NK cells degranulation and cytokine production. We next compared the functionality of inflamed CD56^dim^ NK cells with other NK subsets using the low CD16 and CD226 expression as a proxy to separate them from CD56^bright^ and CD56^dim^ NK cells. We found that CD16/CD226^lo^ CD56^dim^ NK cells had a lower IFNγ and CD107a expression than CD16/CD226^hi^ CD56^dim^ in response to K562 classical NK cell target (**Figure 3I)**. Unlike CD56^bright^ NK cells that also had poor reactivity against K562, CD16/CD226^lo^CD56^dim^ NK cells had a low ability to produce IFNγ in response to IL-12/IL-15/IL-18 stimulation (**Figure 3J)**. Altogether these data reveal the accumulation of CD226/CD16^lo^ NK cell subsets in MM patients with reduced effector functions in response to NKR stimulation.

**Figure 3:**
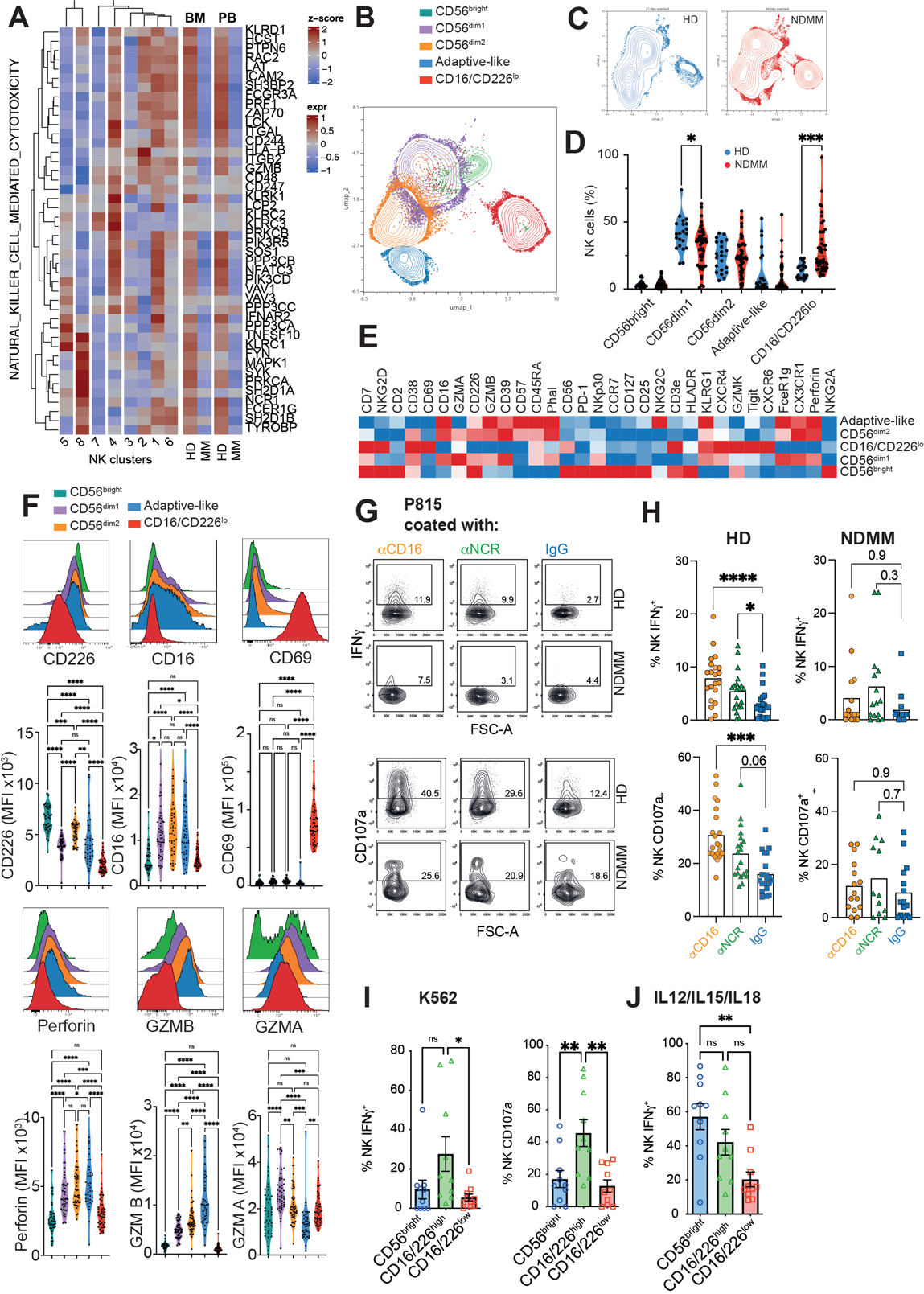
Decreased cytotoxic functions in NK cell from MM patients. **A.** Heatmaps of leading-edge genes standardized expression from “natural_killer_cell_mediated_cytotoxicity” Kegg pathway across NK clusters or disease state (See GSEA analysis Figure 2G-H). **B-F**. A spectral flow cytometry analysis of NK cells from 49 NDMM and 21 HD was performed. **B.** UMAP showing the identification of five main NK subsets by unsupervised clustering. **C-D.** UMAP **(C)** and graphs **(D)** showing differences in cluster distributions among HD and NDMM patients. **(E)** Heatmap showing the expression of the indicated markers by the different NK cell clusters. **(F)** Facs plots and graphs showing the mean fluorescence intensity (MFI) of the indicated markers by the different NK cell clusters. **G.** Purified blood HD and MM NK cells were incubated with P815 coated with anti-CD16, anti-NCR (NKp30, NKp44, NKp46) or control IgG for 6 hrs. Representative Facs plots (**G**) and graphs (**H**) showing the expression of CD107a degranulation marker and the intracellular production of IFNγ. Each dot represents an independent donor. **(I-J)** NK cells purified form MM patients were incubated with K562 (**I**, E:T ratio of 2:1) or in the presence of IL-12/IL-15/IL-18 (**J**) for 6 hrs. Graphs showing the expression of CD107a degranulation marker and the intracellular production of IFNγ by CD56^brigh^, CD16/CD226^low^ and CD16/CD226^high^ CD56^Dim^ NK subsets. Each symbol represents an individual mouse. *p<0.05; **p<0.01, ***p<0.001. ANOVA with Tukey’s post-test analysis.

### Actin remodelling and adhesion defects in NK cells from MM patients

Decreased expression of cell adhesion pathways “Focal adhesion”, “Cell adhesion molecules” and “Regulation of Actin cytoskeleton” were identified as key features of inflamed NK cell clusters C3 and C7 (**Figure 2H, 4A and S4A**), questioning the ability of NK cells from MM patients to interact efficiently with target cells. To address this issue, we first analyzed the interaction between purified NK cells and K562 *in vitro* and observed that NK cells from MM patients had reduced ability to form conjugates with K562 compared to HD (**Figure 4B)**. To better understand the origin of the cell adhesion defects observed in MM NK cells, we focused on the interaction mediated by the β2 integrin LFA-1 (αLβ2) on NK cells with ICAM-1 on target cells. The LFA-1/ICAM-1 interactions lead to actin cytoskeleton remodeling and represents a critical step towards NK cell initial adhesion, immune synapse formation and cytotoxic granule polarization toward target cells ^28^. This involves a switch from LFA-1 inactive bent conformation, to an “extended open” conformation, detectable through the m24 antibody, increasing LFA-1/ICAM-1 affinity ^29^. We investigated differences in immune synapse morphology by confocal microscopy, using MM vs HD NK cell plated on recombinant ICAM-1. Phalloidin staining revealed important differences in polymerized actin organization between HD and MM. HD NK cells formed large asymmetric polymerized actin rings while MM NK cells formed actin ring of lower area and with a less spread shape than HD (**Figure 4 C-D**). We also observed a significant decrease in open LFA-1 spots per synapse in MM NK cells compared to HD (**Figure 4C-D**). Similar results were obtained by comparing FACS sorted CD16/CD226^-^ CD56^dim^ NK with their CD16/CD226^+^ counterparts (**Figure S4C-D)**. Analysis of phalloidin staining by flow cytometry ex vivo indicate that, even at steady state, MM NK cells have lower polymerized actin filaments compared to HD (**Figure 4E)**. Analysis of phalloidin staining among MM NK cells subsets revealed that CD16/CD226^lo^CD56^dim^ MM NK cells had lower polymerized actin content compared to other CD56^dim^ NK subsets (**Figure 4F)**. In accordance with the link between Integrin signaling, actin cytoskeleton remodeling and cellular shape, we also found that CD16/CD226^lo^ NK cell populations had reduced motility after a short exposure to IL-2 than CD16/CD226^hi^ CD56^dim^ counterparts (**Figure 4G-H)**.

**Figure 4:**
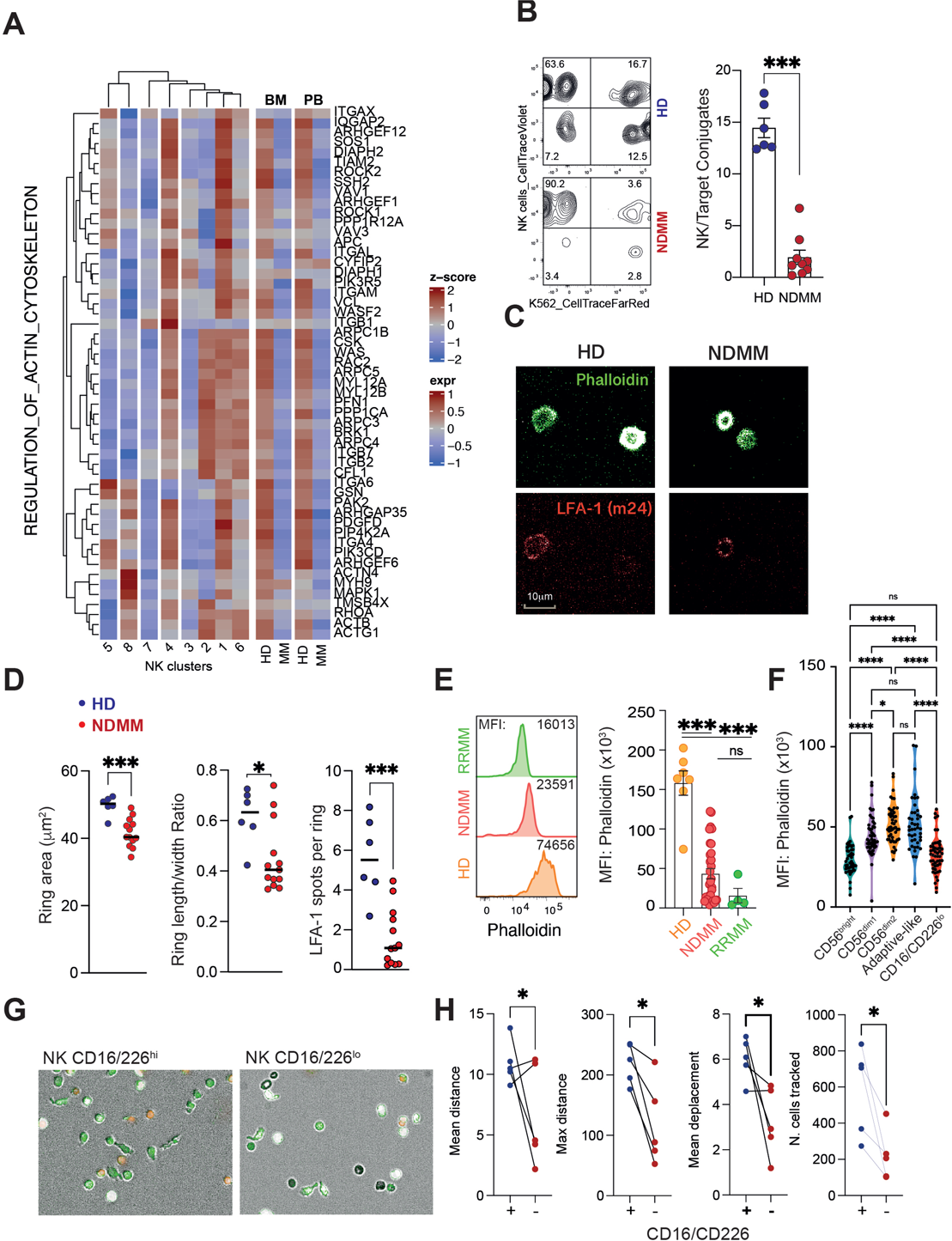
Actin remodelling and adhesion defects in NK cells from MM patients. **A.** Heatmaps of leading-edge genes standardized expression from the “regulation_of_actin_cytoskeleton” Kegg pathway by the indicated NK clusters across NK clusters or disease state (See GSEA analysis Figure 2G-H). **B.** Facs plot and graph showing conjugates between cell trace violet labelled HD or MM NK cells and K562 stained with cell trace far red at a ratio of 1:1 for 1 hour. **C**. Representative pictures showing actin ring formation (phalloidin; green) and open LFA-1 (m24; red) by MM or HD NK cells plated on ICAM-1 coated wells for 20 minutes. **D**. Graph showing the actin ring area, length/width ratio and LFA-1 spots per rings in the indicated conditions. Each dot represents the mean of at least 50 NK cells from each donor. **E**. Phalloidin expression by PB Lin^-^CD56^+^ NK cells from HD, NDMM and RRMM analysed by fow cytometry. **F.** Phalloidin expression by the indicated NK cells subsets from NDMM identified as in Figure 3B. **G-H.** The motility of the indicated NK cell populations labelled with CFSE (green) and PI (Orange) in response to IL-2 was analysed. **G.** Representative pictures. **H.** Graph showing the indicated parameters calculated using the harmony software. Each dot represents the mean of at least 50 NK cells from one donor. Representative histograms and graph showing the phalloidin staining of actin filaments from HD or MM patients NK cells. *p<0.05; **p<0.01, ***p<0.001 Mann Whitney test or ANOVA with Tukey’s post-test analysis.

### Accumulation of altered NK cells in MM patients correlates with poor clinical outcome

Despite the importance of NK cells in MM immunosurveillance and therapy, very little information relates NK cells to clinical outcome ^30^. To answer this question, we analyzed NK cell frequency and phenotype within the BM of a retrospective cohort of 177 MM patients from the IFM 2009 clinical trial involving patients receiving induction therapy with RVD (lenalidomide, bortezomib, dexamethasone) and autologous transplantation ^31^ (**Figure 5A**). We observed an increase in the BM frequency of CD3^-^CD56^+^ NK cells in MM patients as compared to HD (**Figure 5B**). Surprisingly, patients with a high frequency of NK cell (NK^high^ > median) had a lower overall survival (OS) than patients with a low frequency of NK cell (NK^low^::: median; **Figure 5C**). We next analyzed the clinical impact CD16/CD226^neg^ NK cell subsets in the same cohort given their increase and functional alterations in MM patients (**Figure 5D)**. A high frequency of CD226/CD16^neg^ NK cells (CD16/CD226^neg^ > median) had a lower OS than patients with a low frequency of NK cell (CD16/CD226^neg^::: median; **Figure 5D-E).** Of note, other NK cell markers such as CD28, TIGIT, PD-1, CD57, KLRG1, CD38 didn’t have a significant effect on MM patient’s OS (**Figure S5A)**. The frequency of BM NK cells or CD16/CD226^neg^ NK cells did not show any correlation with age, sex or International Staging System (ISS; **Figure S5B-C).** A small increase in the frequency of CD16/CD226^neg^ NK cells was observed in patients bearing Del17p and/or t(4;14) chromosomal alterations (**Figure S5C)** suggesting a potential link between these high risk tumor cytogenic markers (**Figure S5D)** and NK cell status. Interestingly, NK cell frequency was not correlated with CD16/CD226 expression (**Figure 5F)** and the combination of both variables, the absolute CD16/CD226^neg^ NK cells frequency among BM cells, had a strong prognostic value in the IFM 2009 cohort (**Figure 5G and Figure S5E)**. Altogether our results suggest that a high frequency of NK cells with low expression of activation receptors CD16 and CD226 is associated with a poor OS in MM patients.

**Figure 5:**
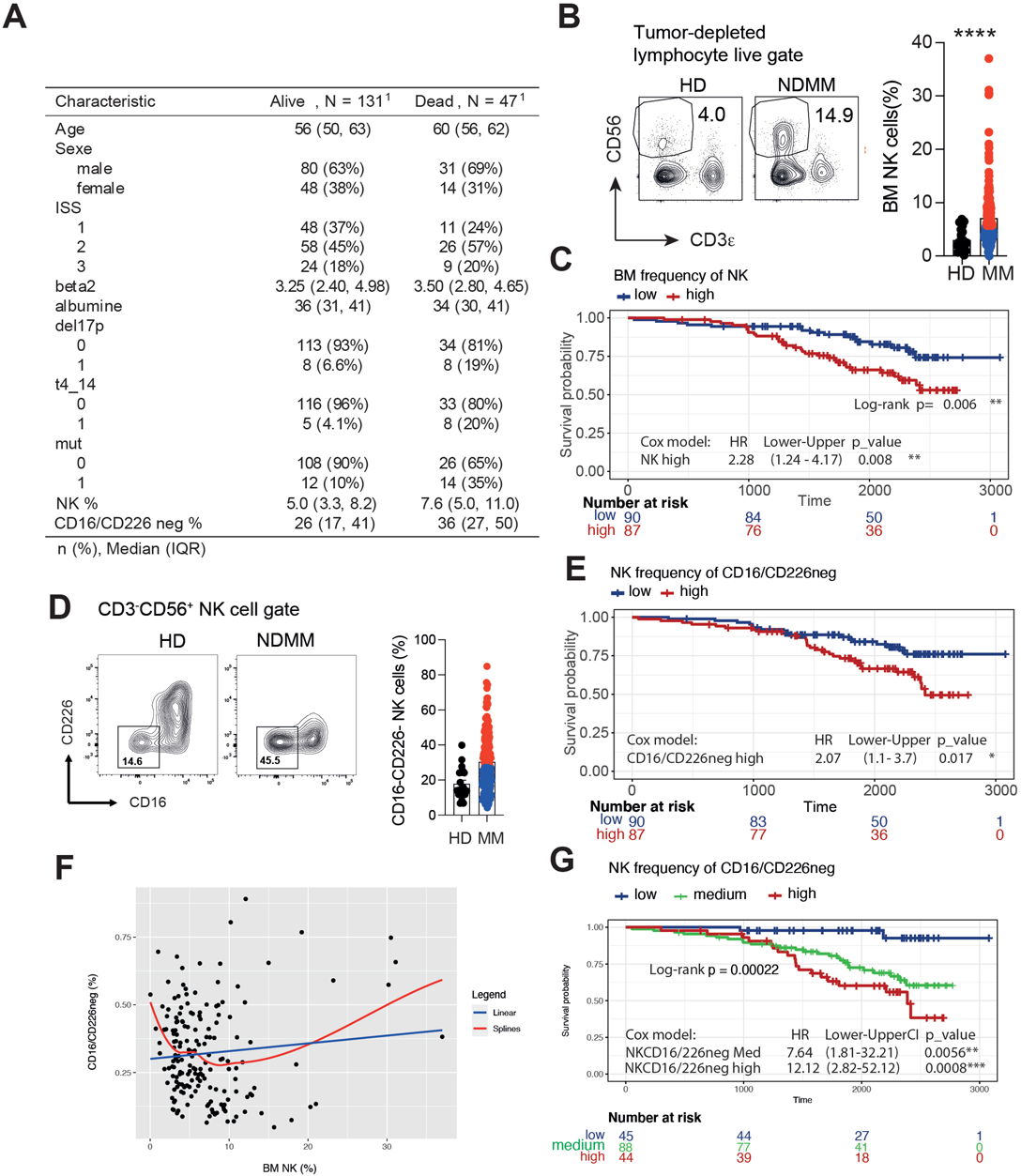
Increased frequency of dysfunctional NK cells in MM patients correlates with poor clinical outcome. NK cells were analyzed by flow cytometry in BM samples of 177 NDMM patients from the IFM2009 clinical trial and 20 HD. **A.** Table showing the clinical features of IFM 2009 myeloma patients involved in this study. **B.** Representative FACS plots and Graph comparing NK cell frequency among CD138-depleted BM from HD and MM patients. **C**. Kaplan–Meier survival estimates over more than 3000 days of follow-up for NK^high^ (>median value) and NK^low^ (:::median value) MM patients. **D**. Representative FACS plots and Graph comparing CD16/CD226negative cell frequency among NK cells from MM patients and HD. **E**. Kaplan– Meier survival estimates over more than 3000 days of follow-up for CD226/CD16neg^high^ (>median value) and CD226/CD16neg^low^ (:::median value) MM patients. **F.** Correlation between BM NK cell and CD16/CD226^negative^ NK cell frequency for the 177 IFM 2009 MM patients. **G.** Kaplan–Meier survival estimates for the absolute frequency of NKCD226/CD16neg among CD138-depleted BM mononuclear cells in IFM 2009 cohort split by quartile.

## DISCUSSION

NK cells are critical immune effector cells for MM immunosurveillance and therapy which call for a better understanding of their landscape in MM patients. In this study, through single cell RNAseq, flow cytometry and functional analysis, we found that MM development is associated with a reduction in Mature cytotoxic “CD56dim” NK cells at the expense of late-stage NK cells characterized by a low CD16 and CD226 expression, inflammatory signatures, adhesion defects and reduced effector functions. The accumulation of these CD16/CD226^lo^ NK cells was associated with a negative clinical outcome in NDMM patients. Given the growing interest in harnessing NK cells to treat myeloma, this improved knowledge around MM-associated NK cell dysfunction will benefit the development of more efficient immunotherapeutic drugs against MM.

By their natural cytotoxic functions and their secretion of proinflammatory cytokines, NK cells are involved in multiple processes of tumor control ^32^. In addition, their safety profile makes these cells attractive targets for cancer immunotherapy. Many strategies are being currently explored to exploit NK cells for cancer treatment with promising preclinical efficacy achieved by chimeric antigen receptor (CAR)-NK cells and NK cell engagers bispecific antibodies in myeloma ^19–21^. Yet, NK-cell-based therapies are limited due to the incomplete understanding of human NK cell heterogeneity and their potential changes within the tumor microenvironment. Improved knowledge about the heterogeneity and distribution of human NK cell subsets emerged from scRNA-seq studies in healthy donors ^22^, but it is still unclear to what extent NK cell transcriptomic landscape from myeloma patients differed from HD. Pan-cancer scRNAseq studies also provided great opportunities to appreciate the large spectrum of tumor-infiltrating NK cell subsets ^33^, still the generalization of the findings are limited by the tumor-type-specific heterogeneity in NK cell composition. We addressed these issues, by performing a scRNAseq analysis of paired blood and BM purified NK cells from NDMM and age/sex matched HD. We found that MM is accompanied by important NK cell alterations. In particular, we observed, in MM patients, a significant decrease in mature cytotoxic CD56^dim^ NK cell clusters at the profit of NK cell clusters characterized by inflammatory signatures and decreased cytotoxic features. Despite their low cytotoxic potential and their low CD16 and CD226 expression, NK cell subsets that accumulated in MM patients had distinct gene signatures and phenotype from CD56^bright^ immature NK cells. Interestingly, trajectory analysis suggested that MM enriched CD16/CD226^lo^ “inflamed” NK cell subsets are late-stage or chronically activated NK cells that would derive from mature CD56^dim^ subsets.

While MM cells reside almost exclusively in the BM compartment, our study revealed similar NK cell alterations in the PB and the BM compartment. Although the mechanism underlying this NK cell phenotypic shifts in MM patients remains unclear, a possible explanation could be the release of myeloma-induced factors into the bloodstream that could affect NK cells at the systemic level. Indeed, MM is characterized by a strong inflammatory network ^34^ and several cytokines detected at high level in patients’ serum could account for NK cell transcriptomic changes observed in MM patients. These include notably NFκB signalling cytokines such as IL-1β and TNF that were recently involved in driving stromal cell inflammation ^34^ or IL-18 that was involved in myeloid derived suppressor cells (MDSC) suppressive functions in MM ^10^. We also found a MM associated NK cell cluster characterized by a set of interferon-stimulated genes (ISGs) suggesting that additional MM associated inflammatory factors, such as type I interferons, may affect NK cell in MM ^35^. It is interesting to underline that many of the aforementioned factors are involved in the expansion of immunosuppressive cells such as MSCs, MDSCs and Tregs within the MM microenvironment ^10,34,35^, that may directly attenuate NK cell activity. This calls for a better understanding of the relative impact of the different MM associated inflammatory mediators on pro-versus anti-tumor immune components.

Like previous studies ^8,36,37^, we found an increased NK cell frequency as compared to HD. Surprisingly, a higher frequency of NK cells was associated with a decrease in OS in MM patients from the IFM2009 cohort. The origin of this result remains poorly understood. While there is possible increase in NK cell frequency with age ^38^, our data didn’t show any correlation between BM NK cell frequency and the age of MM patients. No correlations between NK frequency and other relevant clinical features were identified. Finally, the frequency of NK cells was not linked to their CD16/CD226^lo^ status, but the combination of both parameters (e.g the BM CD16/CD226^lo^ NK cell frequency) had a higher impact on clinical outcome.

The treatment given to MM patients from the IFM2009 clinical trial (e.g Velcade, Revlimid, Dexamethasone and melphalan) didn’t contain specific immunotherapeutic drugs. Still, PI and IMIDs have well described immunomodulatory properties especially on NK cells that could explain the prognostic value of NK cell parameters ^39–42^. The anti-CD38 mAbs, such as daratumumab and isatuximab, in combination with glucocorticoids, PI and Imids now represent the standard-of-care for the treatment of newly diagnosed MM. Evaluating NK cell alterations may be even more relevant with this treatment given the importance of CD16 and NK cell−mediated ADCC in the efficacy of anti-CD38 mAbs ^16,17,43,44^.

## ACKNOWLEDGEMENT

We are grateful to the genotoul bioinformatics platform Toulouse Midi-Pyrenees. This work was granted access to the HPC resources of CALMIP supercomputing center under the allocation P19043. We thank Manon Farcé, Laetitia Ligat and the members from the CRCT core facility. This study has been supported by the Riney Fundation, “la Fondation ARC” (PGA1-20160203788, 20190208630 and SIGN’IT2021), the “Institut National du Cancer” (INCA; PLBIO R16100BB, R19-045, R20-229), by Aviesan PNP grant, by the “French National Research Agency” (ANR LABEX Toucan and the EUR CARe N°ANR-18-EURE-0003), the Fondation Toulouse Cancer Santé, the IUCT-O translational research program and the Intergroupe Francophone du Myélome (IFM). E.B was funded by “la Fondation ARC”.

## AUTHORSHIP CONTRIBUTIONS

Study conception and design: L.M. Acquisition of data: E.B., B.R., M-V.J., M.V., C.M., N.C., M.C., V.B., Analysis and interpretation of data: E.B., B.R., R.E., V.B., H.D., P-P.A., L.M. Drafting of manuscript: P-P.A., L.M. Critical revision and editing: T.W., L.E.L., A.P., J.C., H.A-L., P-P.A., and L.M. Provision of key materials: A.P., J.C., H.A-L.

## REFERENCES

1. Lokhorst HM, Plesner T, Laubach JP, et al. Targeting CD38 with Daratumumab Monotherapy in Multiple Myeloma. N Engl J Med. 2015;373(13):1207–1219.

2. Moreau P, Attal M, Hulin C, et al. Bortezomib, thalidomide, and dexamethasone with or without daratumumab before and after autologous stem-cell transplantation for newly diagnosed multiple myeloma (CASSIOPEIA): a randomised, open-label, phase 3 study. Lancet. 2019;394(10192):29–38.

3. Moreau P, Garfall AL, van de Donk N, et al. Teclistamab in Relapsed or Refractory Multiple Myeloma. N Engl J Med. 2022;387(6):495–505.

4. San-Miguel J, Dhakal B, Yong K, et al. Cilta-cel or Standard Care in Lenalidomide-Refractory Multiple Myeloma. N Engl J Med. 2023;389(4):335–347.

5. Lannes R, Samur M, Perrot A, et al. In Multiple Myeloma, High-Risk Secondary Genetic Events Observed at Relapse Are Present From Diagnosis in Tiny, Undetectable Subclonal Populations. J Clin Oncol. 2022:JCO2101987.

6. Perrot A, Lauwers-Cances V, Tournay E, et al. Development and Validation of a Cytogenetic Prognostic Index Predicting Survival in Multiple Myeloma. J Clin Oncol. 2019;37(19):1657–1665.

7. Guillerey C, Ferrari de Andrade L, Vuckovic S, et al. Immunosurveillance and therapy of multiple myeloma are CD226 dependent. J Clin Invest. 2015;125(7):2904.

8. Larrayoz M, Garcia-Barchino MJ, Celay J, et al. Preclinical models for prediction of immunotherapy outcomes and immune evasion mechanisms in genetically heterogeneous multiple myeloma. Nat Med. 2023;29(3):632–645.

9. Nakamura K, Smyth MJ, Martinet L. Cancer immunoediting and immune dysregulation in multiple myeloma. Blood. 2020;136(24):2731–2740.

10. Nakamura K, Kassem S, Cleynen A, et al. Dysregulated IL-18 Is a Key Driver of Immunosuppression and a Possible Therapeutic Target in the Multiple Myeloma Microenvironment. Cancer Cell. 2018;33(4):634–648 e635.

11. Guillerey C, Harjunpaa H, Carrie N, et al. TIGIT immune checkpoint blockade restores CD8(+) T-cell immunity against multiple myeloma. Blood. 2018;132(16):1689–1694.

12. Weulersse M, Asrir A, Pichler AC, et al. Eomes-Dependent Loss of the Co-activating Receptor CD226 Restrains CD8(+) T Cell Anti-tumor Functions and Limits the Efficacy of Cancer Immunotherapy. Immunity. 2020;53(4):824–839 e810.

13. Carbone E, Neri P, Mesuraca M, et al. HLA class I, NKG2D, and natural cytotoxicity receptors regulate multiple myeloma cell recognition by natural killer cells. Blood. 2005;105(1):251–258.

14. El-Sherbiny YM, Meade JL, Holmes TD, et al. The requirement for DNAM-1, NKG2D, and NKp46 in the natural killer cell-mediated killing of myeloma cells. Cancer Res. 2007;67(18):8444–8449.

15. Kassem S, Diallo BK, El-Murr N, et al. SAR442085, a novel anti-CD38 antibody with enhanced antitumor activity against multiple myeloma. Blood. 2022;139(8):1160–1176.

16. Nijhof IS, Lammerts van Bueren JJ, van Kessel B, et al. Daratumumab-mediated lysis of primary multiple myeloma cells is enhanced in combination with the human anti-KIR antibody IPH2102 and lenalidomide. Haematologica. 2015;100(2):263–268.

17. Viola D, Dona A, Caserta E, et al. Daratumumab induces mechanisms of immune activation through CD38+ NK cell targeting. Leukemia. 2020.

18. Fehniger TA, Cooper MA. Harnessing NK Cell Memory for Cancer Immunotherapy. Trends Immunol. 2016;37(12):877–888.

19. Cichocki F, Bjordahl R, Goodridge JP, et al. Quadruple gene-engineered natural killer cells enable multi-antigen targeting for durable antitumor activity against multiple myeloma. Nat Commun. 2022;13(1):7341.

20. Giang KA, Boxaspen T, Diao Y, et al. Affibody-based hBCMA x CD16 dual engagers for NK cell-mediated killing of multiple myeloma cells. N Biotechnol. 2023;77:139–148.

21. Gauthier L, Morel A, Anceriz N, et al. Multifunctional Natural Killer Cell Engagers Targeting NKp46 Trigger Protective Tumor Immunity. Cell. 2019;177(7):1701–1713 e1716.

22. Crinier A, Milpied P, Escaliere B, et al. High-Dimensional Single-Cell Analysis Identifies Organ-Specific Signatures and Conserved NK Cell Subsets in Humans and Mice. Immunity. 2018;49(5):971–986 e975.

23. Yang C, Siebert JR, Burns R, et al. Heterogeneity of human bone marrow and blood natural killer cells defined by single-cell transcriptome. Nat Commun. 2019;10(1):3931.

24. Melsen JE, Lugthart G, Vervat C, et al. Human Bone Marrow-Resident Natural Killer Cells Have a Unique Transcriptional Profile and Resemble Resident Memory CD8(+) T Cells. Front Immunol. 2018;9:1829.

25. Freud AG, Caligiuri MA. Human natural killer cell development. Immunol Rev. 2006;214:56–72.

26. Hideshima T, Chauhan D, Schlossman R, Richardson P, Anderson KC. The role of tumor necrosis factor alpha in the pathophysiology of human multiple myeloma: therapeutic applications. Oncogene. 2001;20(33):4519–4527.

27. Costes V, Portier M, Lu ZY, Rossi JF, Bataille R, Klein B. Interleukin-1 in multiple myeloma: producer cells and their role in the control of IL-6 production. Br J Haematol. 1998;103(4):1152–1160.

28. Gross CC, Brzostowski JA, Liu D, Long EO. Tethering of intercellular adhesion molecule on target cells is required for LFA-1-dependent NK cell adhesion and granule polarization. J Immunol. 2010;185(5):2918–2926.

29. Comrie WA, Babich A, Burkhardt JK. F-actin flow drives affinity maturation and spatial organization of LFA-1 at the immunological synapse. J Cell Biol. 2015;208(4):475–491.

30. Orrantia A, Terren I, Astarloa-Pando G, et al. NK Cell Reconstitution After Autologous Hematopoietic Stem Cell Transplantation: Association Between NK Cell Maturation Stage and Outcome in Multiple Myeloma. Front Immunol. 2021;12:748207.

31. Attal M, Lauwers-Cances V, Hulin C, et al. Lenalidomide, Bortezomib, and Dexamethasone with Transplantation for Myeloma. N Engl J Med. 2017;376(14):1311–1320.

32. Martinet L, Smyth MJ. Balancing natural killer cell activation through paired receptors. Nat Rev Immunol. 2015;15(4):243–254.

33. Tang F, Li J, Qi L, et al. A pan-cancer single-cell panorama of human natural killer cells. Cell. 2023;186(19):4235–4251 e4220.

34. de Jong MME, Kellermayer Z, Papazian N, et al. The multiple myeloma microenvironment is defined by an inflammatory stromal cell landscape. Nat Immunol. 2021;22(6):769–780.

35. Kawano Y, Zavidij O, Park J, et al. Blocking IFNAR1 inhibits multiple myeloma-driven Treg expansion and immunosuppression. J Clin Invest. 2018;128(6):2487–2499.

36. Frassanito MA, Silvestris F, Cafforio P, Silvestris N, Dammacco F. IgG M-components in active myeloma patients induce a down-regulation of natural killer cell activity. Int J Clin Lab Res. 1997;27(1):48–54.

37. Zavidij O, Haradhvala NJ, Mouhieddine TH, et al. Single-cell RNA sequencing reveals compromised immune microenvironment in precursor stages of multiple myeloma. Nat Cancer. 2020;1(5):493–506.

38. Brauning A, Rae M, Zhu G, et al. Aging of the Immune System: Focus on Natural Killer Cells Phenotype and Functions. Cells. 2022;11(6).

39. Soriani A, Zingoni A, Cerboni C, et al. ATM-ATR-dependent up-regulation of DNAM-1 and NKG2D ligands on multiple myeloma cells by therapeutic agents results in enhanced NK-cell susceptibility and is associated with a senescent phenotype. Blood. 2009;113(15):3503–3511.

40. Shi J, Tricot GJ, Garg TK, et al. Bortezomib down-regulates the cell-surface expression of HLA class I and enhances natural killer cell-mediated lysis of myeloma. Blood. 2008;111(3):1309–1317.

41. Lundqvist A, Yokoyama H, Smith A, Berg M, Childs R. Bortezomib treatment and regulatory T-cell depletion enhance the antitumor effects of adoptively infused NK cells. Blood. 2009;113(24):6120–6127.

42. Lundqvist A, Berg M, Smith A, Childs RW. Bortezomib Treatment to Potentiate the Anti-tumor Immunity of Ex-vivo Expanded Adoptively Infused Autologous Natural Killer Cells. J Cancer. 2011;2:383–385.

43. Moreno L, Perez C, Zabaleta A, et al. The Mechanism of Action of the Anti-CD38 Monoclonal Antibody Isatuximab in Multiple Myeloma. Clin Cancer Res. 2019;25(10):3176–3187.

44. Verkleij CPM, Frerichs KA, Broekmans MEC, et al. NK Cell Phenotype Is Associated With Response and Resistance to Daratumumab in Relapsed/Refractory Multiple Myeloma. Hemasphere. 2023;7(5):e881.

