## Supplemental material for "Inflamed Natural Killer cells with adhesion defects are associated with a poor prognosis in Multiple Myeloma"

**SUPPLEMENTARY METHODS:**

**Single cell RNA sequencing**

Pre-processing and quality control

Alignment and quantitation (including intronic reads) were performed with the “cellranger count” command for each emulsion (*cellranger* v.6.1.2 using the human 10x genomics reference) to generate unique molecular identifier (UMI) RNA and antibody derived tags (ADT) count matrices. Additional spliced RNA and unspliced RNA UMI count matrices were computed using *Velocyto* [59] and used to calculate a nuclear RNA fraction defined as the fraction of intronic counts among total counts. Modelling nuclear fraction as gaussian mixture allowed the identification of damaged vs intact cells (high vs low fractions) as implemented in *DropletQC* [60]. To detect doublets, we trained a classifier on simulated heterogenous doublets and applied it to differentiate single cells from doublets, as implemented in scDblFinder [61].

We then annotated cell types against a public reference PBMC dataset using the *Seurat* R package [62]. We applied the single cell transformation (SCT) to RNA counts. We then mapped cells from our dataset to the reference using the “anchor” algorithm implemented in *Seurat*, yielding per-cell predictions of queried cells against reference cell types.

We performed all above steps on each emulsion separately and then excluded damaged cells, doublets, as well as cells with library size or number of genes detected below 500, library size above 12000 and percentage of counts mapping to mitochondrial chromosome genes above 10%. We also excluded cells with a predicted main immune cell type (‘l1’ granularity) confidence score below 0.9. This quality control ensured high quality immune cells were conserved and we retained only cells that mapped to the NK reference cell type (‘l1’ granularity). Given high inter-emulsion variability of the ADT signal, we did not further consider ADT data for our analysis.

<u>Clustering and dimensionality reduction</u>

For cells passing quality control, we combined cells from all emulsions to form a combined dataset and computed normalized UMI counts by dividing each count by the total number of counts per cell. We then multiplied normalized counts by 10,000 and added a pseudo count of 1 before log-transformation. We computed the stabilized variance of each gene using the variance-stabilizing transformation (VST) and retained the top 2000 hypervariable genes (HVGs) with highest stabilized variance for principal component analysis (PCA). We z-scored HVGs and regressed out percentage of UMI mapping to mitochondrial chromosome genes as well as percentage of UMI mapping to ribosomal genes. We then computed the first 50 principal components (PCs) using a partial singular value decomposition method, based on the implicitly restarted Lanczos bidiagonalization algorithm (IRLBA), as implemented in the *Seurat* R package. To correct for systematic differences across donors, we applied harmony integration [63] to the first 50 PC loadings and retained 22 harmony-corrected PCs to build nearest neighbor graphs for visualization using Uniform Manifold Approximation and Projection (UMAP), and community detection using Leiden algorithm, as implemented in *Seurat* with default parameters.

We identified 9 clusters (resolution parameter = 0.5), including a small proliferating NK cluster which we excluded from our analysis. We then reapplied our dimensionality reduction pipeline to conserved cells, from identifying HVGs, to UMAP and clustering as described above for our final analysis, yielding 8 NK cells clusters.

<u>Marker genes prioritization and GSEA analysis</u>

For each gene, we computed Receiver operating characteristic (ROC) curves between pair of clusters and used area under the curve (AUC) as a ranking metric to prioritize cluster marker genes and run pre-ranked geneset enrichment analysis (GSEA). For each tested geneset, the union of all leading-edge genes across all paired comparisons were used to build custom gene modules and compute enrichment scores at the single cell level: calculated as the average expression of the gene module, subtracted by the average expression of a randomly generated gene module, through the *addModuleScore* function of *Seurat*, using normalized UMI counts. For heatmap visualizations of manually curated genes, log-transformed gene expression was averaged for each category (cluster, patient, disease and tissue) and resulting values z-scored or rescaled between 0 and 1.

Trajectory analysis

Using the harmony-corrected PCs as input, we computed a minimum spanning tree (MST) on clusters centroids and derived pseudotime by building principal curves based on the MST branches (labelled as lineages) as implemented in *slingshot* [64]. We computed the average of pseudotime across all lineages, weighted by the probability of belonging to each lineage, to obtain an overall pseudotime value for each cell and use for all visualizations.
