## Supplemental figures for "Inflamed Natural Killer cells with adhesion defects are associated with a poor prognosis in Multiple Myeloma"

### FIGURE S1

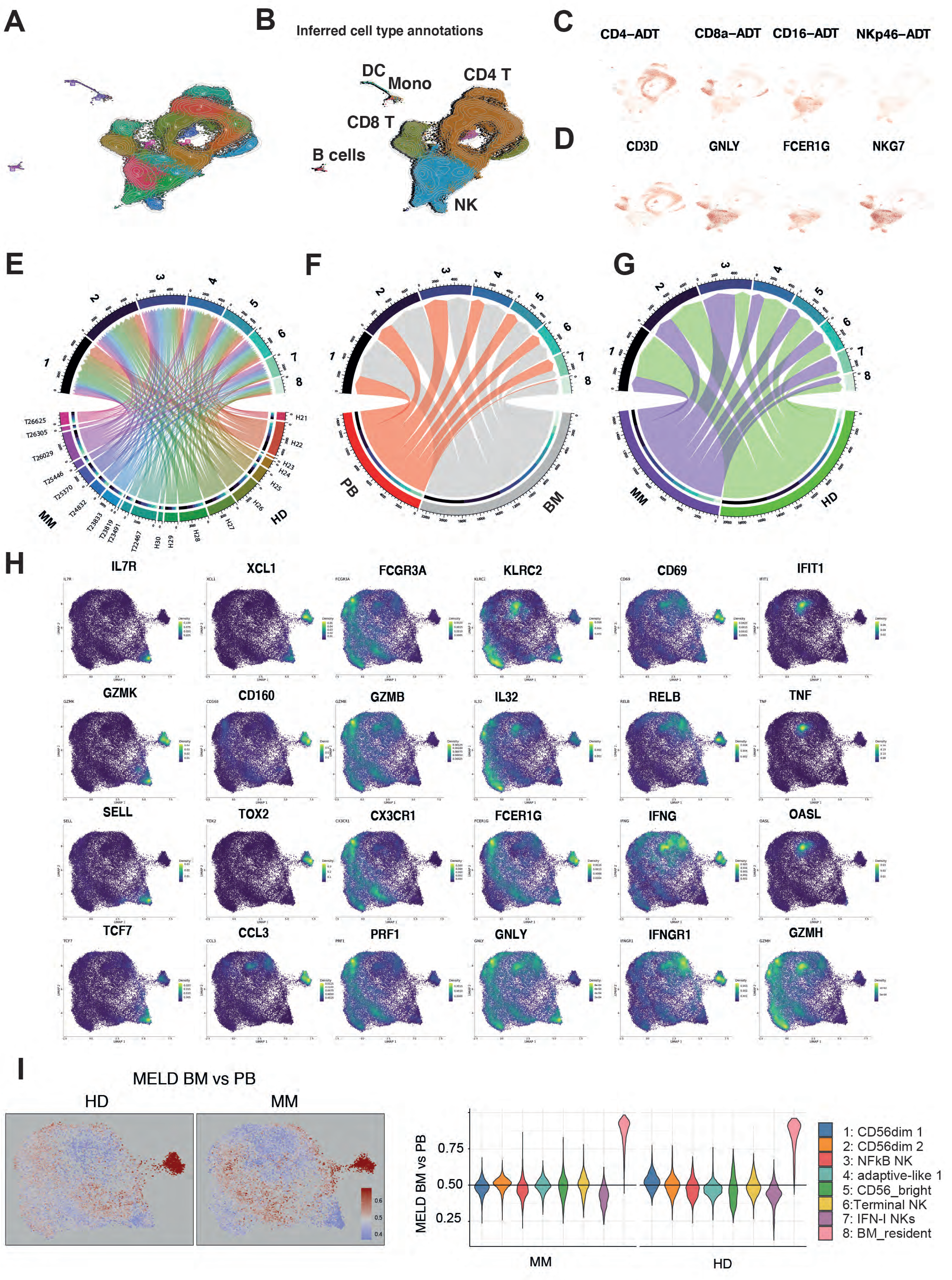
A
B
Inferred cell type annotations
DC
Mono
CD4 T
CD8 T
B cells
NK
C
CD4–ADT
CD8a–ADT
CD16–ADT
NKp46–ADT
D
CD3D
GNLY
FCER1G
NKG7
E
F
G
MM
HD
PB
BM
H
IL7R
XCL1
FCGR3A
KLRC2
CD69
IFIT1
GZMK
CD160
GZMB
IL32
RELB
TNF
SELL
TOX2
CX3CR1
FCER1G
IFNG
OASL
TCF7
CCL3
PRF1
GNLY
IFNGR1
GZMH
I
MELD BM vs PB
HD
MM
MELD BM vs PB
0.75
0.50
0.25
MM
HD
1: CD56dim 1
2: CD56dim 2
3: NFkB NK
4: adaptive-like 1
5: CD56_bright
6:Terminal NK
7: IFN-I NKs
8: BM_resident

**Figure S1: Sc RNA analysis of blood and BM NK and T cells from 10 MM patients and 10 HD.**

**A.** UMAP showing the clustering of 109690 paired blood and BM $CD3^{-}CD56^{+}$ NK and $CD3^{+}$ T cells from 10 MM and 10 age/sex HD. **B.** UMAP showing the predicted immune cell types against a public reference PBMC dataset using the Seurat R package (Hao, Cell, 2021). **C-D.** Expression of the T cell and NK cell surface proteins **(C)** and genes **(D)** visualized on global UMAP. **E-G.** Circos plots showing the relative contribution of each sample (**E**), tissue type (**F**, PB or BM) and condition (**G**, MM or HD) in the 8 NK cell clusters shown in **Figure 1A**. **H.** Relative expression of the indicated genes on global UMAP. **I.** MELD likelihood (see supplementary Methods) highlighting BM vs PB enrichment computed separately on HD or MM. Likelihood values are overlayed on UMAP embeddings and per-cluster likelihood distributions displayed as violin plots.

### FIGURE S2

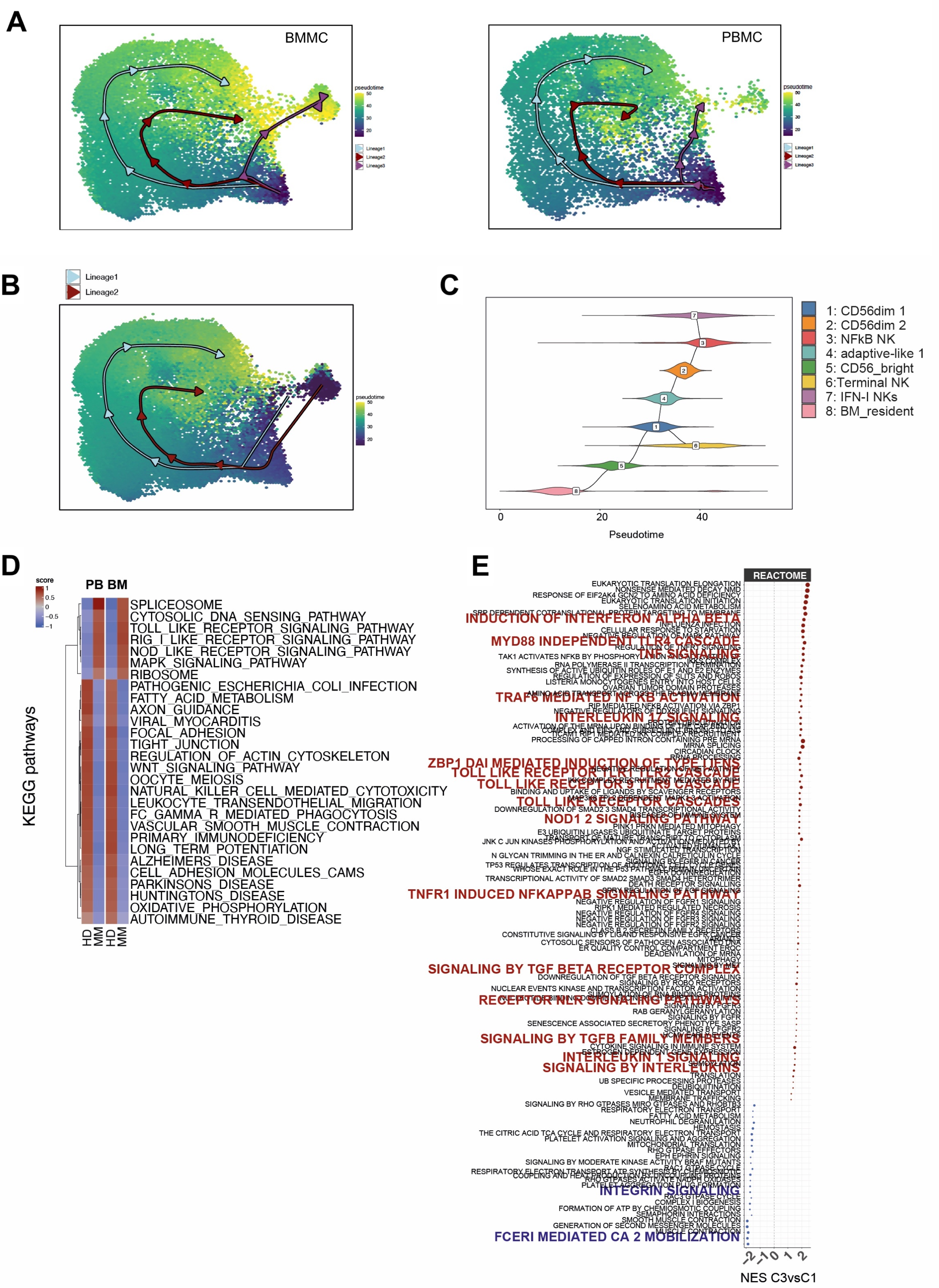

A
BMMC
PBMC
pseudotime
50
40
30
20
Lineage1
Lineage2
Lineage3
B
Lineage1
Lineage2
pseudotime
C
1: CD56dim 1
2: CD56dim 2
3: NFkB NK
4: adaptive-like 1
5: CD56_bright
6:Terminal NK
7: IFN-I NKs
8: BM_resident
0
20
40
Pseudotime
D
score
1
0.5
0
−0.5
−1
PB BM
SPLICEOSOME
CYTOSOLIC_DNA_SENSING_PATHWAY
TOLL_LIKE_RECEPTOR_SIGNALING_PATHWAY
RIG_I_LIKE_RECEPTOR_SIGNALING_PATHWAY
NOD_LIKE_RECEPTOR_SIGNALING_PATHWAY
MAPK_SIGNALING_PATHWAY
RIBOSOME
PATHOGENIC_ESCHERICHIA_COLI_INFECTION
FATTY_ACID_METABOLISM
AXON_GUIDANCE
VIRAL_MYOCARDITIS
FOCAL_ADHESION
TIGHT_JUNCTION
REGULATION_OF_ACTIN_CYTOSKELETON
WNT_SIGNALING_PATHWAY
OOCYTE_MEIOSIS
NATURAL_KILLER_CELL_MEDIATED_CYTOTOXICITY
LEUKOCYTE_TRANSENDOTHELIAL_MIGRATION
FC_GAMMA_R_MEDIATED_PHAGOCYTOSIS
VASCULAR_SMOOTH_MUSCLE_CONTRACTION
PRIMARY_IMMUNODEFICIENCY
LONG_TERM_POTENTIATION
ALZHEIMERS_DISEASE
CELL_ADHESION_MOLECULES_CAMS
PARKINSONS_DISEASE
HUNTINGTONS_DISEASE
OXIDATIVE_PHOSPHORYLATION
AUTOIMMUNE_THYROID_DISEASE
KEGG pathways
HD
MM
HD
MM
E
REACTOME
EUKARYOTIC TRANSLATION ELONGATION
NONSENSE MEDIATED DECAY NMD
RESPONSE OF EIF2AK4 GCN2 TO AMINO ACID DEFICIENCY
EUKARYOTIC TRANSLATION INITIATION
SELENOAMINO ACID METABOLISM
INDUCTION OF INTERFERON ALPHA BETA
INFLUENZA INFECTION
CELLULAR RESPONSE TO STARVATION
MYD88 INDEPENDENT TLR4 CASCADE
TNF SIGNALING
TRAF6 MEDIATED NF KB ACTIVATION
INTERLEUKIN 17 SIGNALING
MRNA SPLICING
CIRCADIAN CLOCK
RRNA PROCESSING
TOLL LIKE RECEPTOR TLR1 TLR2 CASCADE
TOLL LIKE RECEPTOR 9 CASCADE
BINDING AND UPTAKE OF LIGANDS BY SCAVENGER RECEPTORS
TOLL LIKE RECEPTOR CASCADES
NOD1 2 SIGNALING PATHWAY
PINK1 PRKN MEDIATED MITOPHAGY
N GLYCAN TRIMMING IN THE ER AND CALNEXIN CALRETICULIN CYCLE
TRANSCRIPTIONAL ACTIVITY OF SMAD2 SMAD3 SMAD4 HETEROTRIMER
DEATH RECEPTOR SIGNALLING
TNFR1 INDUCED NFKAPPAB SIGNALING PATHWAY
NEGATIVE REGULATION OF FGFR1 SIGNALING
RIPK1 MEDIATED REGULATED NECROSIS
NEGATIVE REGULATION OF FGFR4 SIGNALING
NEGATIVE REGULATION OF FGFR3 SIGNALING
NEGATIVE REGULATION OF FGFR2 SIGNALING
CLASS B 2 SECRETIN FAMILY RECEPTORS
ER QUALITY CONTROL COMPARTMENT ERQC
DEADENYLATION OF MRNA
MITOPHAGY
SIGNALING BY TGF BETA RECEPTOR COMPLEX
DOWNREGULATION OF TGF BETA RECEPTOR SIGNALING
SIGNALING BY ROBO RECEPTORS
NUCLEAR EVENTS KINASE AND TRANSCRIPTION FACTOR ACTIVATION
SUMOYLATION OF RNA BINDING PROTEINS
RECEPTOR NLR SIGNALING PATHWAYS
SIGNALING BY FGFR3
RAB GERANYLGERANYLATION
SIGNALING BY FGFR
SENESCENCE ASSOCIATED SECRETORY PHENOTYPE SASP
SIGNALING BY FGFR2
SIGNALING BY TGFB FAMILY MEMBERS
CYTOKINE SIGNALING IN IMMUNE SYSTEM
ESTROGEN DEPENDENT GENE EXPRESSION
INTERLEUKIN 1 SIGNALING
SUMOYLATION
SIGNALING BY INTERLEUKINS
TRANSLATION
UB SPECIFIC PROCESSING PROTEASES
DEUBIQUITINATION
VESICLE MEDIATED TRANSPORT
MEMBRANE TRAFFICKING
SIGNALING BY RHO GTPASES MIRO GTPASES AND RHOBTB3
RESPIRATORY ELECTRON TRANSPORT
FATTY ACID METABOLISM
NEUTROPHIL DEGRANULATION
HEMOSTASIS
THE CITRIC ACID TCA CYCLE AND RESPIRATORY ELECTRON TRANSPORT
PLATELET ACTIVATION SIGNALING AND AGGREGATION
MITOCHONDRIAL TRANSLATION
RHO GTPASE EFFECTORS
EPH EPHRIN SIGNALING
SIGNALING BY MODERATE KINASE ACTIVITY BRAF MUTANTS
RAC1 GTPASE CYCLE
PLATELET AGGREGATION PLUG FORMATION
INTEGRIN SIGNALING
RAC3 GTPASE CYCLE
COMPLEX I BIOGENESIS
FORMATION OF ATP BY CHEMIOSMOTIC COUPLING
SEMAPHORIN INTERACTIONS
SMOOTH MUSCLE CONTRACTION
GENERATION OF SECOND MESSENGER MOLECULES
MUSCLE CONTRACTION
FCERI MEDIATED CA 2 MOBILIZATION
-2
-1
0
1
2
NES C3vsC1

**Figure S2: Accumulation of late stage inflamed NK cells in Multiple myeloma.**

A. Trajectory analysis run separately on BMMC vs PBMC show no major differences. B. Trajectory analysis results overlayed on UMAP embeddings identified 2 lineages seeding from BMr NK cluster 8. C. Branching of the minimum spanning tree built at the cluster level. D. Heatmaps comparing MM and HD standardized expression of the NK clusters C7vsC4 and C3vsC1 differentially expressed KEGG pathways (See Figure 2H). E. Comparison of gene expression programs between cluster 3 and 1 using GSEA analysis of REACTOME database. Some relevant pathways related to inflammation and Cell adhesion were manually highlighted.

### FIGURE S3

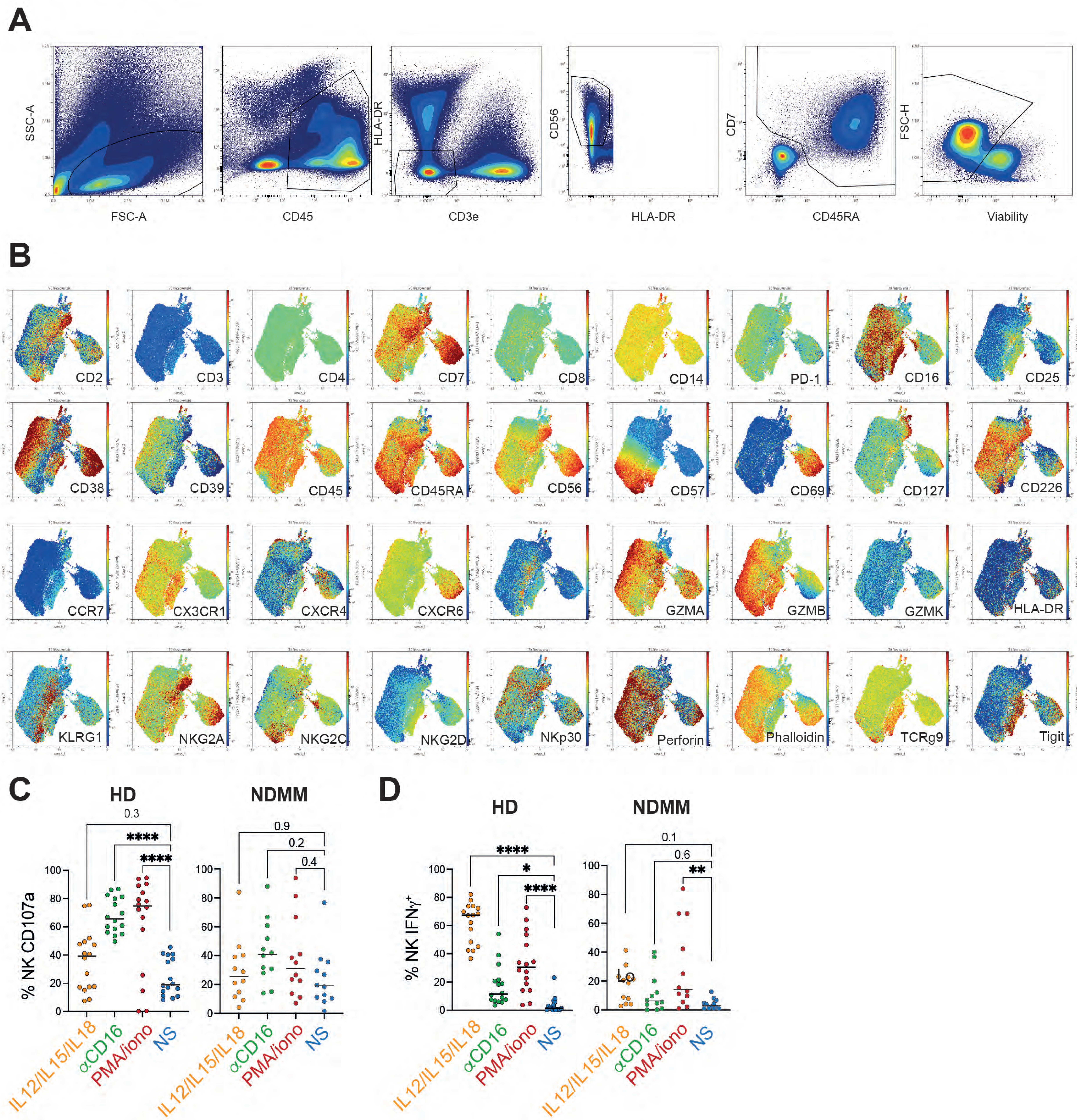
A
SSC-A
FSC-A
CD45
HLA-DR
CD3e
CD56
HLA-DR
CD7
CD45RA
FSC-H
Viability
B
CD2
CD3
CD4
CD7
CD8
CD14
PD-1
CD16
CD25
CD38
CD39
CD45
CD45RA
CD56
CD57
CD69
CD127
CD226
CCR7
CX3CR1
CXCR4
CXCR6
GZMA
GZMB
GZMK
HLA-DR
KLRG1
NKG2A
NKG2C
NKG2D
NKp30
Perforin
Phalloidin
TCRg9
Tigit
C
HD
NDMM
0.3
****
****
0.9
0.2
0.4
% NK CD107a
100
80
60
40
20
0
IL12/IL15/IL18
αCD16
PMA/iono
NS
D
HD
NDMM
****
*
****
0.1
0.6
**
% NK IFNγ+
100
80
60
40
20
0
IL12/IL15/IL18
αCD16
PMA/iono
NS

**Figure S3: Decreased cytotoxic functions in NK cell from MM patients.**

**A-B.** A spectral flow cytometry analysis of NK cells from 49 NDMM and 21 HD was performed. **A.** FACS plots imllustrating the gating strategy used to define NK cells. **B.** UMAP showing the relative expression of the indicated markers by the different NK cell cluster as in **Figure 3B**. **C-D.** Purified blood HD and MM NK cells were plated in wells, coated with Ig control (NS), with anti-CD16 mAbs, in the presence of IL-12/IL-15/IL-18 or PMA/ionomycine for 6 hrs. Graphs showing the expression of CD107a degranulation marker (**C)** and the intracellular production of IFNγ **(D)**. Each dot represents an independent donor. *p<0.05; **p<0.01, ***p<0.001. ANOVA with Tukey's post-test analysis.

FIGURE S4

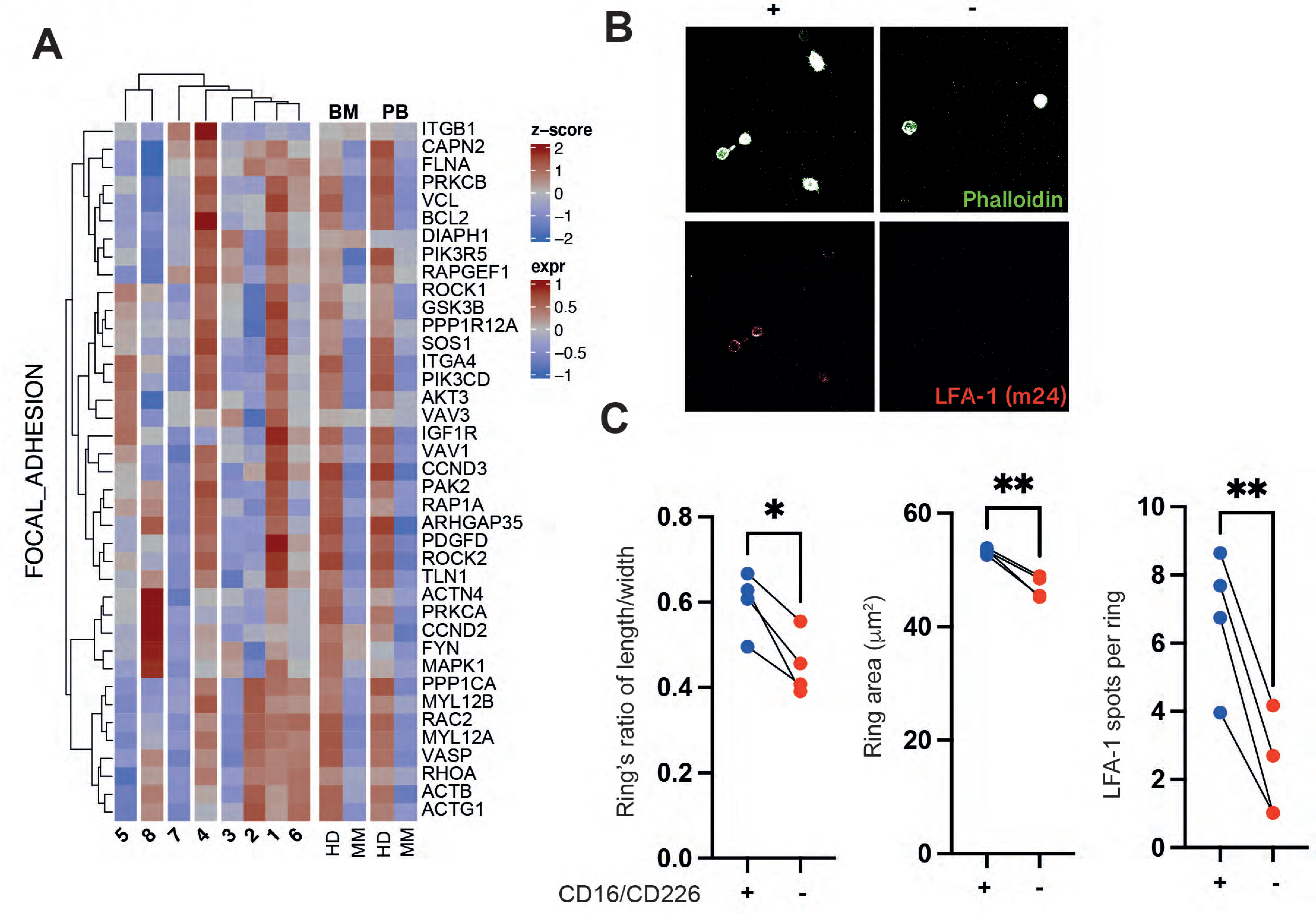
A
FOCAL_ADHESION
BM
PB
z-score
2
1
0
−1
−2
expr
1
0.5
0
−0.5
−1
ITGB1
CAPN2
FLNA
PRKCB
VCL
BCL2
DIAPH1
PIK3R5
RAPGEF1
ROCK1
GSK3B
PPP1R12A
SOS1
ITGA4
PIK3CD
AKT3
VAV3
IGF1R
VAV1
CCND3
PAK2
RAP1A
ARHGAP35
PDGFD
ROCK2
TLN1
ACTN4
PRKCA
CCND2
FYN
MAPK1
PPP1CA
MYL12B
RAC2
MYL12A
VASP
RHOA
ACTB
ACTG1
5
8
7
4
3
2
1
6
HD
MM
HD
MM
B
CD16/CD226
+
-
Phalloidin
LFA-1 (m24)
C
Ring's ratio of length/width
0.8
0.6
0.4
0.2
0.0
*
CD16/CD226
+
-
Ring area (μm²)
60
40
20
0
**
+
-
LFA-1 spots per ring
10
8
6
4
2
0
**
+
-

**Figure S4: Reduced size and adhesion defects in NK cells from MM patients**

**A.** Heatmaps of leading-edge gene standardized expression from the "FOCAL_ADHESION" Kegg pathway across NK clusters or disease state (See GSEA analysis **Figure 2G-H**). **B**. Representative pictures showing actin ring formation (phalloidin; green) and open LFA-1 (m24; red) by sorted CD16/CD226$^{low}$ and CD16/CD226$^{high}$ CD56$^{dim}$ NK cells plated on ICAM1 coated wells for 20 minutes. **C**. Graph showing the actin ring area, length/width ratio and LFA-1 spots per rings in the indicated conditions. Each dot represents the mean of at least 50 NK cells from each donor. *p<0.05; **p<0.01, ***p<0.001 Paired T-test or ANOVA with Tukey's post-test analysis.

### FIGURE S5

## A

| Characteristic | HR | p |
|---|---|---|
| CD28+ (%) | 1.2033997 | 0.14943331 |
| TIGIT+ (%) | 0.9352809 | 0.66749989 |
| PD1+ (%) | 0.8046353 | 0.40638745 |
| CD57+ (%) | 0.8245876 | 0.19017907 |
| KLRG1+ (%) | 0.9053161 | 0.53208302 |
| CD38+ (%) | 1.2201885 | 0.25181014 |

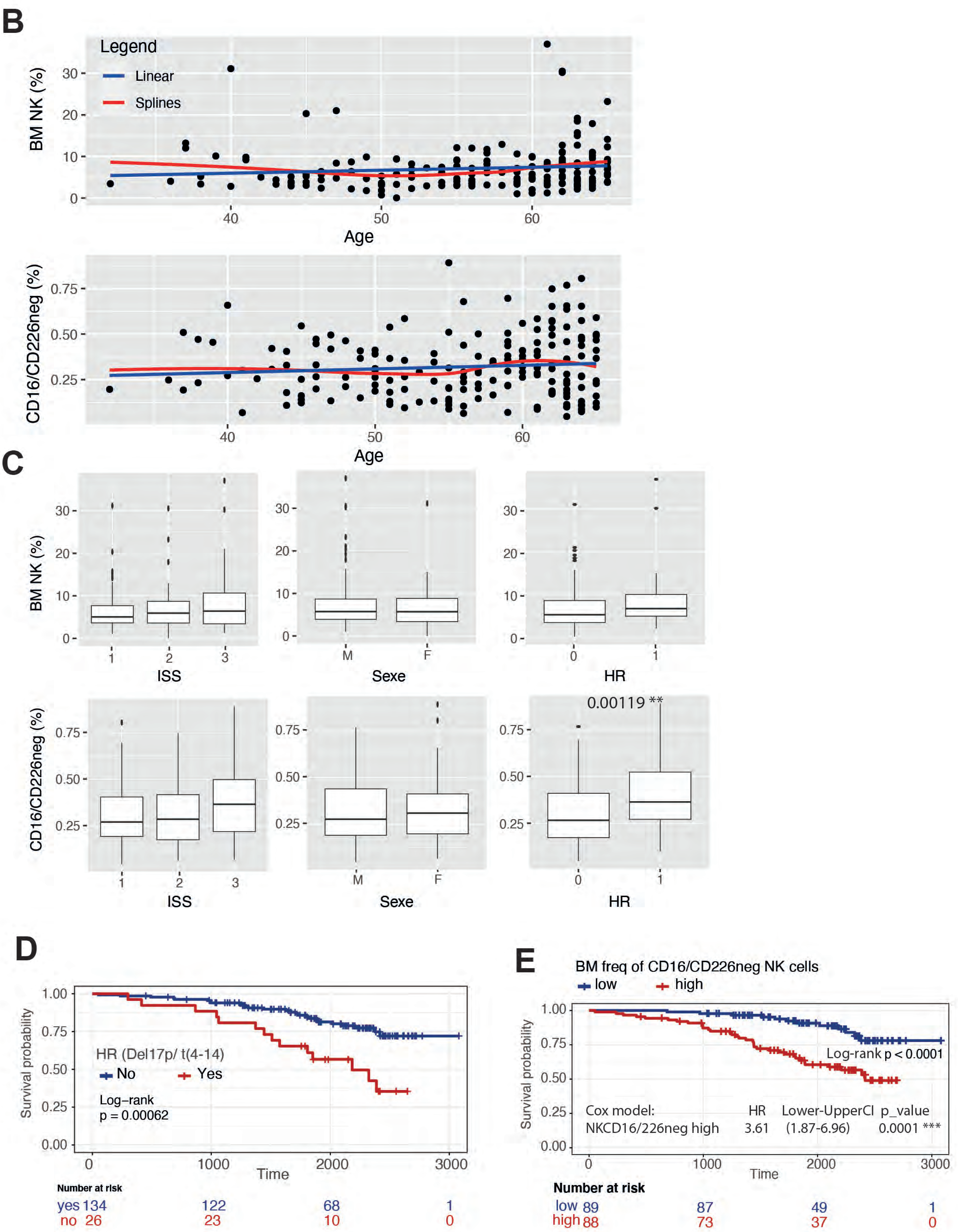

**Figure S5: Increased frequency of dysfunctional NK cells in MM patients correlates with poor clinical outcome.**

**A.** Table showing the survival HR and p value of the indicated NK cell parameters in a linear cox model. **B.** Graph showing the correlation between the age of the patients and the frequency of BM NK cell or CD16/CD226negative NK cell for the 177 patients analysed in the IFM 2009 cohort. **C.** box-and-whisker plots showing the BM NK cell or CD16/CD226negative NK cell frequencies according to the ISS, gender or (Del17p and/or T4:14) mutational status of the 177 patients analysed in IFM 2009 cohort. **D**. Kaplan–Meier survival estimates for patients with or without highr risk tumor alterations (Del17p and/or T4:14). **E.** Kaplan–Meier survival estimates for the absolute frequency of NKCD226/CD16neg among CD138-depleted BM mononuclear cells in IFM 2009 cohort split by median.
